# Human hip osteoarthritis–associated *Chadl* variant induces intervertebral disc degeneration in mice

**DOI:** 10.64898/2026.06.04.730159

**Authors:** Kimheak Sao, Bryn Mallon, Jacqueline Shine, Anastasia Mavridis, Elena V. Filippova, Jack R. Felkner, Carson D. Shepler, Tanner M. Orders, Susan D’Costa, William A. Cowan, Brian O. Diekman, Makarand V. Risbud

## Abstract

Chondroadherin-like (CHADL), a small leucine-rich proteoglycan, plays a role in ECM assembly and chondrocyte differentiation. CHADL has been implicated in osteoarthritis (OA) due to the discovery of a rare frameshift variant (rs532464664) that confers one of the highest genetic risks for hip OA reported to date. In this study, we characterized the skeletal phenotypes of two mouse models of *Chadl* loss-of-function, a global null and an 8bp insertion that mimics the human frameshift variant. Surprisingly, while there was a slight increase in knee OA at 12 months, there were no genotype-based differences in knee or hip OA in aged mice. Interestingly, histological assessment revealed phenotypic changes in the intervertebral discs of mutant mice, along with increased disc height index, suggesting altered motion-segment biomechanics. Polarized imaging showed dysregulated collagen turnover in the annulus fibrosus (AF), evidenced by changes in the proportions of thin and intermediate fibers. There was decreased abundance of the nucleus pulposus (NP) marker CA3 and decreased staining for ECM proteins in the AF of mutants. CHADL loss affected vertebral trabecular architecture, cortical thickness, and mineral density. This study highlights an unrecognized role of CHADL in fine-tuning ECM homeostasis and health of the intervertebral disc and vertebral bone.

## INTRODUCTION

Osteoarthritis (OA) and intervertebral disc degeneration are highly prevalent musculoskeletal disorders and leading contributors to pain, disability, and health care utilization worldwide (1, 2). Although OA has classically been investigated in synovial joints such as the knee and hip, degenerative changes in the intervertebral disc share substantial overlap in molecular pathways, biomechanical susceptibilities, and extracellular matrix (ECM) dysregulation (3, 4). Increasing evidence indicates that common genetic determinants may influence disease susceptibility across distinct skeletal tissues (5, 6). Dissecting how individual genetic risk factors give rise to tissue-specific degenerative outcomes remains a fundamental challenge in musculoskeletal biology.

OA is a multifactorial condition arising from complex interactions among aging, mechanical loading, inflammation, and genetic predisposition (7). Genome-wide association studies have identified numerous loci linked to OA risk; however, the causal genes and underlying mechanisms of these loci are only beginning to be investigated (8-12). Among recently identified OA-associated genes, chondroadherin-like (*CHADL*) has emerged as a particularly compelling candidate (13). When present in a homozygous state, a rare frameshift variant in *CHADL* confers one of the largest genetic effect sizes reported for hip OA, highlighting a potentially central role for this gene in maintaining skeletal tissue homeostasis (13, 14). Despite this strong genetic association, the *in vivo* consequences of *Chadl* loss-of-function and its roles beyond articular cartilage degeneration remain largely unexplored.

*CHADL* encodes a non-glycanated, small leucine-rich proteoglycan (SLRP) belonging to the chondroadherin family, a group of ECM-associated proteins known to regulate collagen fibrillogenesis, matrix organization, and cell-matrix interactions (15). Members of this family are critical for the structural integrity and biomechanical properties of connective tissues, including cartilage, tendon, and intervertebral disc (16). Prior *in vitro* and developmental studies implicate CHADL in chondrocyte differentiation and ECM assembly, suggesting that it may function as a modulator of tissue-specific matrix properties (15). However, direct experimental evidence linking CHADL to the maintenance of adult skeletal tissues has been lacking.

The intervertebral disc, the largest avascular and hypoxic tissue, is a highly specialized fibrocartilaginous structure that permits spinal flexibility while withstanding complex mechanical loads (6, 17). Disc degeneration is characterized by progressive ECM breakdown, altered collagen organization, cell death, and loss of phenotypic markers that define the identity of the nucleus pulposus (NP) and annulus fibrosus (AF) (18, 19). While genetic influences on intervertebral disc degeneration are substantial, relatively few genes have been mechanistically linked to disc-specific degeneration (20-23). Notably, a recent multi-ancestry meta-analysis of genome-wide association studies (GWAS) has linked up to 87 loci to chronic back pain, although their link to disc degeneration has not been established (24). Given its established role in ECM regulation, CHADL is an appealing candidate for influencing disc integrity; however, whether CHADL dysfunction preferentially impacts the disc relative to other skeletal tissues remains to be determined.

Human genetic data provides a compelling rationale for addressing this question experimentally. The identification of a high-risk *CHADL* frameshift variant associated with hip OA raises the hypothesis that CHADL loss-of-function may broadly compromise joint and skeletal tissue health (13). At the same time, growing evidence underscores the tissue specificity of OA risk alleles, with certain variants exerting pronounced effects in select joints while sparing others (9). Defining the tissue specificity of CHADL-dependent pathology is therefore critical for interpreting OA genetic risk and elucidating disease mechanisms.

To address these knowledge gaps, we employed complementary mouse models of *Chadl* loss-of-function, including a traditional null allele (deleting a large region of a critical exon) and an 8–base-pair insertion mutation analogous to the human OA-associated variant (13). These models enabled a systematic, age-dependent evaluation of skeletal phenotypes across multiple tissues. We demonstrate that CHADL deficiency results in a selective vulnerability of the lumbar intervertebral discs, characterized by increased disc height, disrupted collagen organization within the AF, and loss of key ECM and NP phenotypic markers. Polarized light imaging further revealed shifts in collagen fibril thickness distributions, consistent with impaired collagen turnover and matrix maturation. In contrast to expectations based on human hip OA genetics, CHADL loss did not produce substantial OA pathology in the knee or hip, underscoring the importance of tissue-specific genetic effects. Instead, our findings identify CHADL as a previously unrecognized regulator of intervertebral disc ECM homeostasis rather than a global determinant of articular cartilage integrity. Additionally, CHADL deficiency altered vertebral bone architecture, affecting trabecular organization, cortical thickness, and mineral density, further supporting a role for CHADL in spine biology.

Collectively, these findings redefine the functional landscape of CHADL in the adult skeleton and identify the intervertebral disc as a primary tissue vulnerable to CHADL dysfunction. By integrating human genetic insights with *in vivo* modeling, this work advances understanding of how specific ECM regulators drive tissue-selective degeneration and underscores the need to consider disc pathology when interpreting OA genetic risk. Such insights are essential for developing targeted therapeutic strategies to prevent or slow the progression of spine degeneration.

## RESULTS

### *Chadl* loss has a subtle effect on OA in articular joints

In order to investigate functions of CHADL in skeletal tissues as a function of age, we generated two complementary mouse models of CHADL loss-of-function (Fig. 1A): a) mice with a 897 base pair deletion in a critical exon of *Chadl* (KO), b) mice harboring an 8-bp insertion mutation in the *Chadl* gene (8bp) resulting in a frameshift, premature termination codon, and nonsense mediated decay of the transcript. Sanger sequencing of the founder 8bp mice and progeny showed the specific 8 base pair insertion that matches the identified human mutation with no other changes to the surrounding sequence (Fig. 1B). Cartilage was obtained from the femoral head of 12-day old mice for Western blot to confirm that deletion of CHADL protein was achieved. Both KO and 8 bp lacked CHADL protein but showed strong bands for GAPDH (housekeeping) and COL6a1 (to confirm the presence of pericellular matrix in the protein preparation) (Fig. 1C).

**Figure 1:**
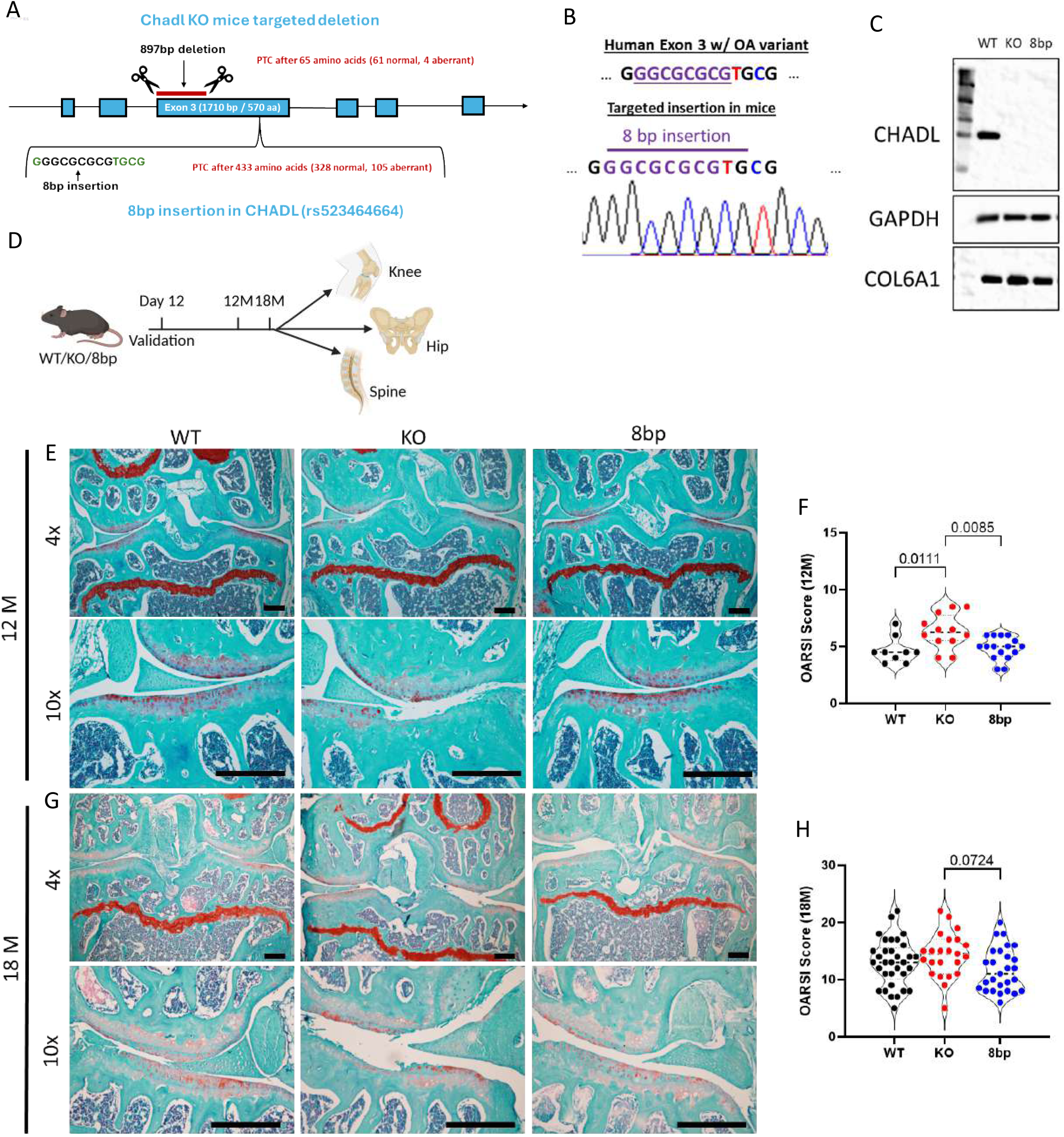
Knee osteoarthritis in mice with *Chadl* loss. (A) Schematic for the generation of *Chadl* KO and 8bp mice. (B) Sanger sequencing showing the insertion of the 8 base pair sequence that mimics the human mutation into the analogous region of the mouse genome. (C) Western blot for Chadl in the hip cartilage of 2-week-old mice, with GAPDH and COLVI as the housekeeping proteins, representing cellular and extracellular matrix (ECM) markers. (D) Schematic showing the experimental design of the study. (E) Safranin O/Fast Green/Hematoxylin staining of paraffin-embedded knee (stifle) joint sections in 12-month-old mice. 10x magnification shows the lateral side. (F) Total joint OARSI score (sum of 4 quadrants with a maximum score of 6 per quadrant) for 12-month-old mice. (G) Representative images of knee joints from 18-month-old mice. (H) OARSI score for aged mice (≥ 18 months). Scale bar for histology images 200 µm. For 12M: 9 WT, 12 KO, and 16 *Chadl* 8bp female mice were used. For aged mice: 33 WT (23F, 10M); 23 KO (10F, 13M), 17 *Chadl* 8bp mice (10F, 7M) were used.

To assess the impact of CHADL loss, these mice were aged and analyzed at 12 months (middle-aged) and 18 months (old) for their skeletal phenotypes (Fig. 1D). Since CHADL has been closely linked to chondrocyte differentiation and the human variant has a strong association with OA (13-15), we first evaluated hindlimb OA using the stifle (knee) as there are standard histological scoring systems for this joint (25). At 12 months, KO mice showed minor to moderate OA changes, such as loss of GAG staining in the matrix and surface fibrillation (Fig. 1E). The severity of OA was determined using a total joint OARSI score. There was a significant effect of genotype (ANOVA main effect p=0.0035), with *Chadl* KO mice having significantly higher total joint OARSI score as compared to wild type (p=0.0101) and *Chadl* 8 bp (p=0.0078) mice (Fig. 1F). When we analyzed older mice, more severe OA-like changes became apparent; however, OARSI scores were comparable between WT and both *Chadl* mutants (Fig. 1G, H). In this aged cohort, genotype showed a trend for an effect on OA, but this effect was not statistically significant (ANOVA main effect, p=0.0788). The strongest trend was a higher OARSI score in *Chadl* KO mice compared with the *Chadl* 8 bp mice (Fig. 1H: 14.26 ± 0.8081 vs. 11.7 ± 0.7310, p=0.0632). While less commonly investigated in mice, we also sectioned and evaluated the hip joint because the human variant was discovered in the context of hip OA. There was no evidence of substantial degeneration of hip joint in old mice of any genotype (Fig. S1), indicating that this joint is more protected than the knee joint. Overall, the *Chadl* KO mice showed only a moderate acceleration of early OA changes in the knee, and the joints from *Chadl* 8 bp mice were no different from wild-type joints.

### CHADL loss promotes intervertebral disc degeneration

Since CHADL is a collagen-interacting SLRP that contributes to cellular mechanosensing and matrix homeostasis (15), which are critical for intervertebral disc function, we investigated whether this protein contributes to intervertebral disc health. To this end, we examined lumbar discs from 12-month-old and 18-month-old mice and performed histopathological assessment of disc tissue morphology using the modified Thompson grading scheme. Safranin O/Fast Green/Hematoxylin staining of the lumbar disc sections of 12-month-old mice did not show significant alterations in NP and AF tissue morphology in *Chadl* mutant lines compared to WT mice (Fig. 2A). On the other hand, 18-month-old *Chadl* mutant mice showed significant signs of disc degeneration. Notably, degeneration was characterized by fewer vacuolated NP cells, morphological changes in inner AF cells, diminished demarcation between the NP and AF compartments, lamellar buckling, and occasional clefts in the outer AF (Fig. 2B). Accordingly, while NP and AF compartment degeneration scores at 12 months were comparable across all genotypes, at 18 months they were significantly higher in both KO and 8bp mice than in wild-type mice (Fig. 2C, D). Notably, more than 50% of AF in 18-month old KO mice showed severe degeneration and were consistently scored grade 3 or higher, in contrast to WT AF tissue with a grade score of 1 or 2 (Fig. 2E). Furthermore, we performed micro-computed tomography (μCT) analysis to determine the changes in disc height and disc height index in experimental mice (Fig. 2F). At 12 months, *Chadl* mutant mice exhibited a pronounced increase in disc height and disc height index (DHI), which are early indicators of disc degeneration, implying altered disc mechanical behavior (Fig. 2G-I). At 18 months, the 8 bp mutant continued to exhibit increased disc height and DHI compared to WT mice. These findings suggest that loss of CHADL function makes mice prone to age-dependent disc degeneration.

**Figure 2.**
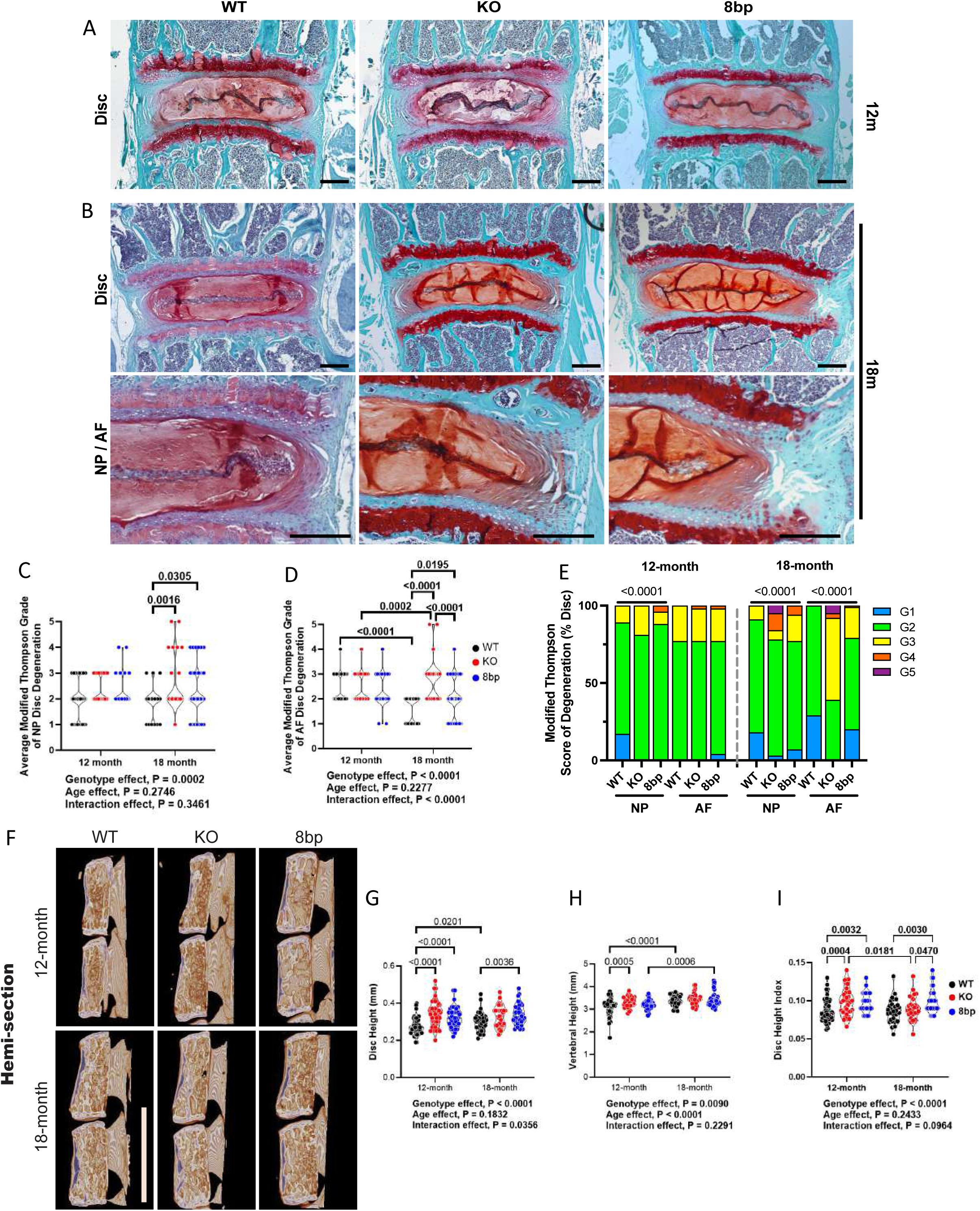
Mice with *Chadl* loss show increased intervertebral disc degeneration with aging. (A) Safranin O/Fast Green/Hematoxylin staining of paraffin embedded disc sections from WT, KO, and 8bp mice at 12M and (B) 18M. Including high magnification image showing degenerative features and loss of NP and AF compartment demarcation at 18 months. (C) Modified Thompson Grading of Lumbar discs in WT, KO, and 8bp mice at 12M and (D) 18M. (E) Distribution of Modified Thompson Grading Scores in 12M and 18Mmice mice. (F) Representative μCT images showing a hemi section of spine in lateral plane used for disc height index measurements. Scale bar = 2.5 mm (G) Vertebral Height (mm), (H) Disc Height (mm), and (I) Disc Height Index measurements. Violin plots show score distribution with median and quartile range, *P* < 0.05. For 12M: 12 WT mice (8F, 4M), n = 65 discs; 10 KO mice (6F, 4M), n = 47 discs; 8 *Chadl* 8bp mice (6F, 2M), n = 48 discs. For 18M: 6 WT (3F, 3M), n=34 discs; 6 KO (1F, 5M), n = 36 discs; 19 *Chadl* 8bp (6F, 13M), n =101 discs. L1/2-6/S1 levels were included in the analysis. Significance was determined using 2-way ANOVA with Tukey’s Comparison comparing Age and Gene Interaction (B, C, F-H) and Chi-square test (D). scale bar for histology images 250 µm.

### CHADL loss alters collagen fiber composition of the disc

To examine collagen fiber structure and changes in thickness accompanying disc degeneration observed in *Chadl* mutants, we performed picrosirius red staining with polarized light microscopy. Notably, at 12 months, 8bp *Chadl* mutant showed an increase and decrease in the fraction of thin and intermediate collagen fibers in the AF compartment compared to WT, respectively (Fig. 3A, B). There were, however, no statistical differences in total fiber fractions across genotypes (Fig. 3B, C). At 18 months, both 8bp mutant and KO mice showed a significant decrease in intermediate fiber fraction compared to WT mice, with no noticeable changes in the thin or thick fibers (Fig.3D-F). These results clearly indicated increased collagen turnover early on, which persisted as mice aged, but without a compensatory increase in thin fibers. While changes in collagen turnover were evident in the AF, there were no corresponding changes in the NP compartment, and all genotypes showed a comparable incidence of NP tissue fibrosis, as observed with disc aging (Fig. 3G).

**Figure 3.**
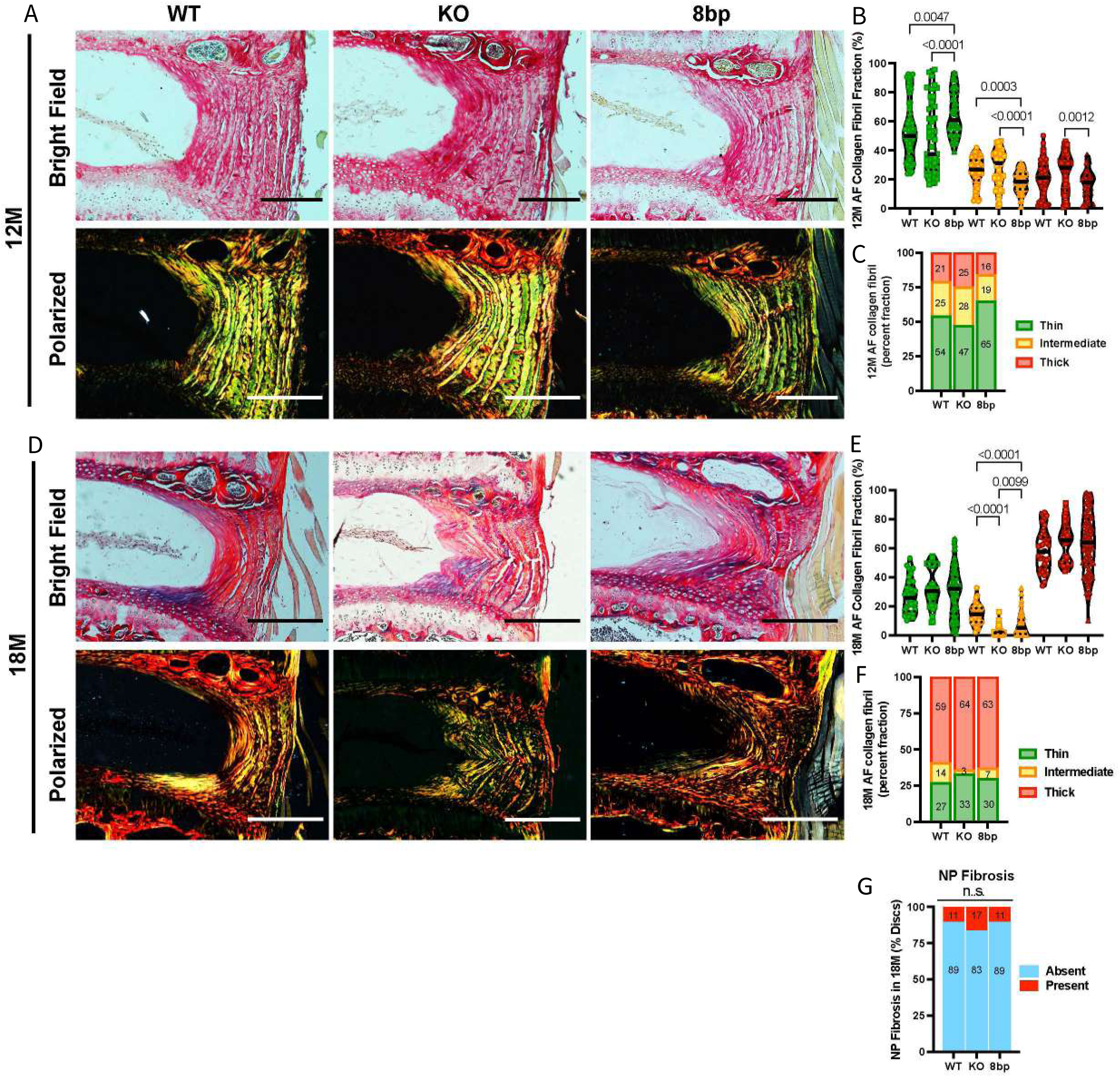
*Chadl* loss changes disc collagen fiber composition. (A) Representative images showing Picro-Sirius Red stained WT, KO, 8bp discs at 12M. (B) AF Collagen Fibril Fraction Measurements at 12M. (C) Distribution of AF collagen fibril fraction at 12M. (D) Representative images showing Picro-Sirius Red stained WT, KO, 8bp discs at 18M. at 18M. (E) AF Collagen Fibril Fraction Measurements at 18M. (F) Distribution of AF collagen fibril fraction at 18M. (G) Percent of discs with NP fibrosis in 18M animals. Violin plots show score distribution with median and quartile range, *P* < 0.05. For 12M: WT mice (8F, 4M), n =70 discs; 10 KO (6F, 4M), n = 48 discs; 8 Chadl 8bp (6F, 2M), n = 49 discs. For 18M: 6 WT (3F, 3M), n = 36 discs; 6 KO (1F, 5M), n=35 discs; 19 *Chadl* 8bp (6F, 13M), n = 89 discs. Significance was determined using 2-way ANOVA with Tukey’s Post Hoc Test comparing Age and Gene interaction (B, D) and Chi square test (P<0.05 significance) (C, F, G). Scale bar 250µm.

### CHADL helps maintain healthy disc ECM and cell phenotype

Given that CHADL loss results in disc degeneration and alters collagen fiber composition, we studied the localization and abundance of select disc ECM molecules by immunohistochemistry. We noted a significant decrease in the abundance of collagen I (COL I) in the AF compartment of KO and 8bp mutants compared to WT discs at 18 months, with no discernible genotype-based difference at 12 months (Fig. 4A, B), suggesting that the observed changes in fiber composition primarily affected COL I fibrils. Notably, collagen II (COL II) staining revealed a similar pattern of lower abundance in the AF of KO than WT mice at 18 months, with a trend of decreasing levels in 8bp mutant; again, there were no differences in COL II abundance at 12 months (Fig. 4C, D). Interestingly, in contrast to collagens, cartilage oligomatrix protein (COMP) staining in the AF compartment of KO and 8bp mutants showed a significant decrease in abundance at 12 months, with no changes at 18 months, suggesting an earlier destabilizing effect on COMP in CHADL mutants (Fig. 5A, B). We also determined the abundance of fibromodulin (FMOD), an important constituent of AF ECM, and observed a decrease in 8bp mice, with a similar trend in KO mice at 12 months. FMOD levels at 18 months were comparable across genotypes, suggesting that aging is a dominant modulator of FMOD levels (Fig. 5C, D). Next, we determined the effect of CHADL loss on aggrecan (ACAN), a major ECM constituent of NP tissue. As expected, ACAN levels were significantly affected by aging, and all genotypes showed an age-dependent decline in abundance without a notable genotype-based difference (Fig. 6A, B). We determined whether the NP cell phenotype is affected in CHADL mutants by measuring the levels of the key marker, CA3 (Fig. 6C, D). Notably, while NP cells preserved their phenotype and showed comparable CA3 levels in WT mice with aging, CA3 abundance decreased in KO and 8bp mice. Moreover, at 18 months, compared to WT mice, CA3 staining showed lower abundance in 8bp mutant, with a trend of decreasing levels in KO (Fig. 6C, D). Overall, our results showed that loss of CHADL dysregulates collagens and other key constituents of disc ECM and affects NP cell phenotype.

**Figure 4.**
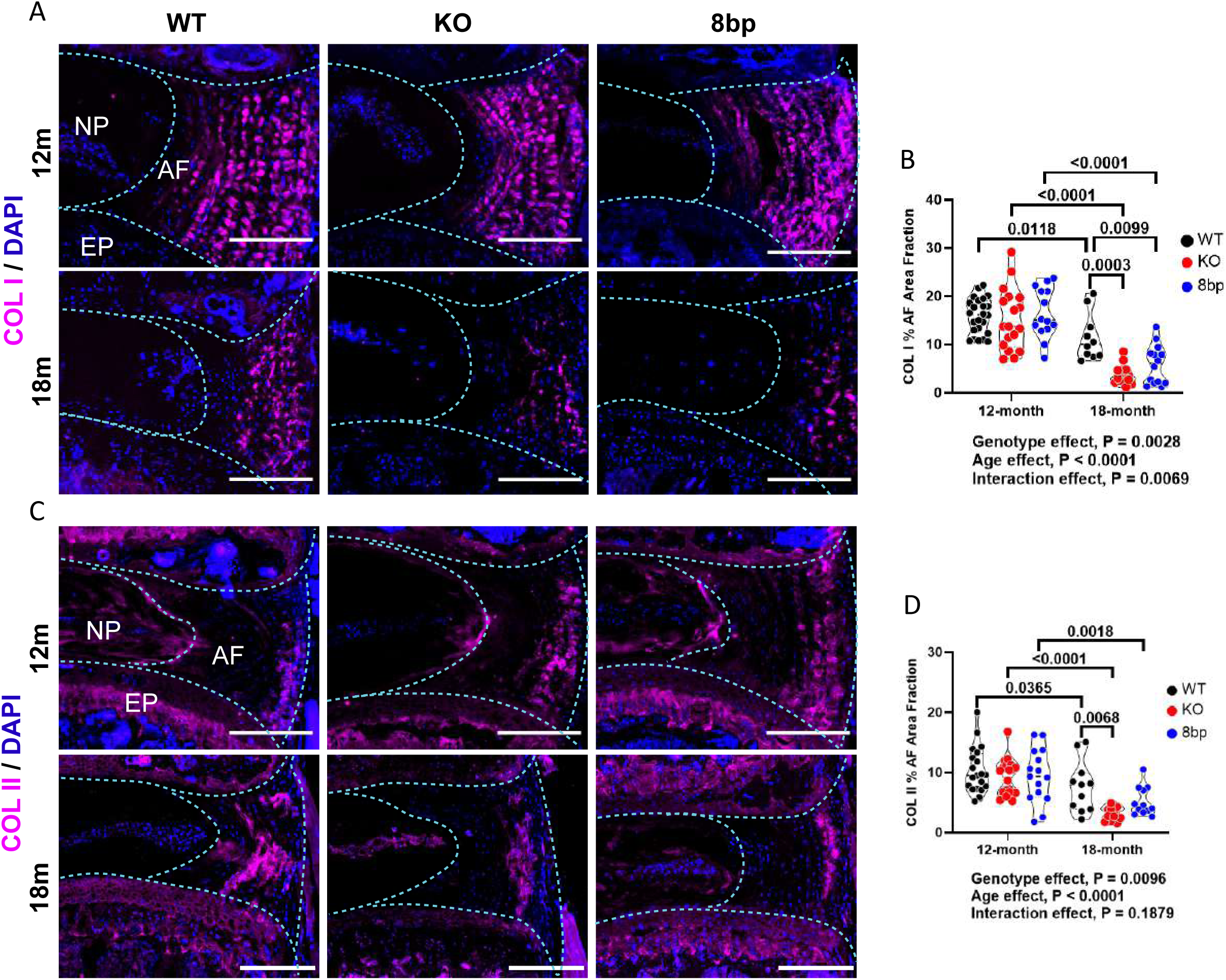
*Chadl* loss causes diminished disc abundance of Collagens I and II. (A) Representative images showing COL I staining of 12M and 18M disc sections. (B) Percent area quantification of COLI in the AF compartment. (C) COLII staining of 12M and 18M disc sections, (D) Percent area quantification of COLII in AF compartment. Violin plots show score distribution with median and quartile range, *P* < 0.05. For 12M: 5-8 WT, KO, 8bp mice, n = 2-4 discs/mouse, 14-24 discs total per genotype. For 18M: 3-6 WT, KO, 8bp mice, n = 2-4 discs/mouse, 9-14 discs total per genotype. Significance was determined using a 2-way ANOVA with Tukey’s Post Hoc Test, comparing Age and Gene interaction. Scale bar 250µm.

**Figure 5.**
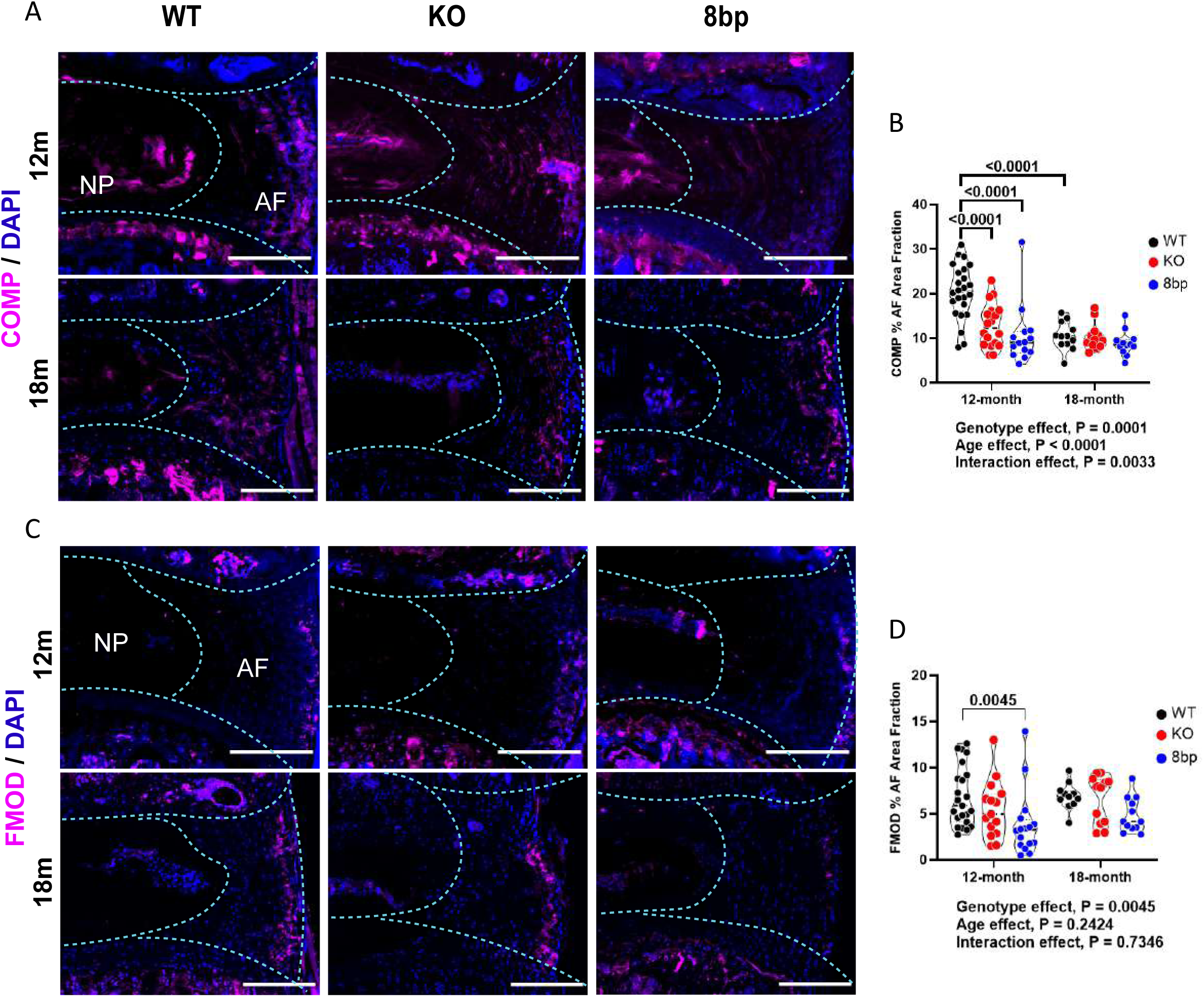
Chadl loss alters disc ECM composition. (A) Representative images showing COMP staining of 12M and 18M disc sections. (B) Percent area quantification of COMP in AF compartment. (C) Fibromodulin staining of 12M and 18M disc sections. (D) Percent area quantification of FMOD in the AF compartment. Violin plots show score distribution with median and quartile range, *P* < 0.05. For 12M: 5-8 WT, KO, 8bp mice, n = 2-4 discs/mouse, 14-24 discs total per genotype. For 18M: 3-6 WT, KO, 8bp mice, n = 2-4 discs/mouse, 9-14 discs total per genotype. Significance was determined using a 2-way ANOVA with Tukey’s Post Hoc Test, comparing Age and Gene interaction. Scale bar 250µm.

**Figure 6.**
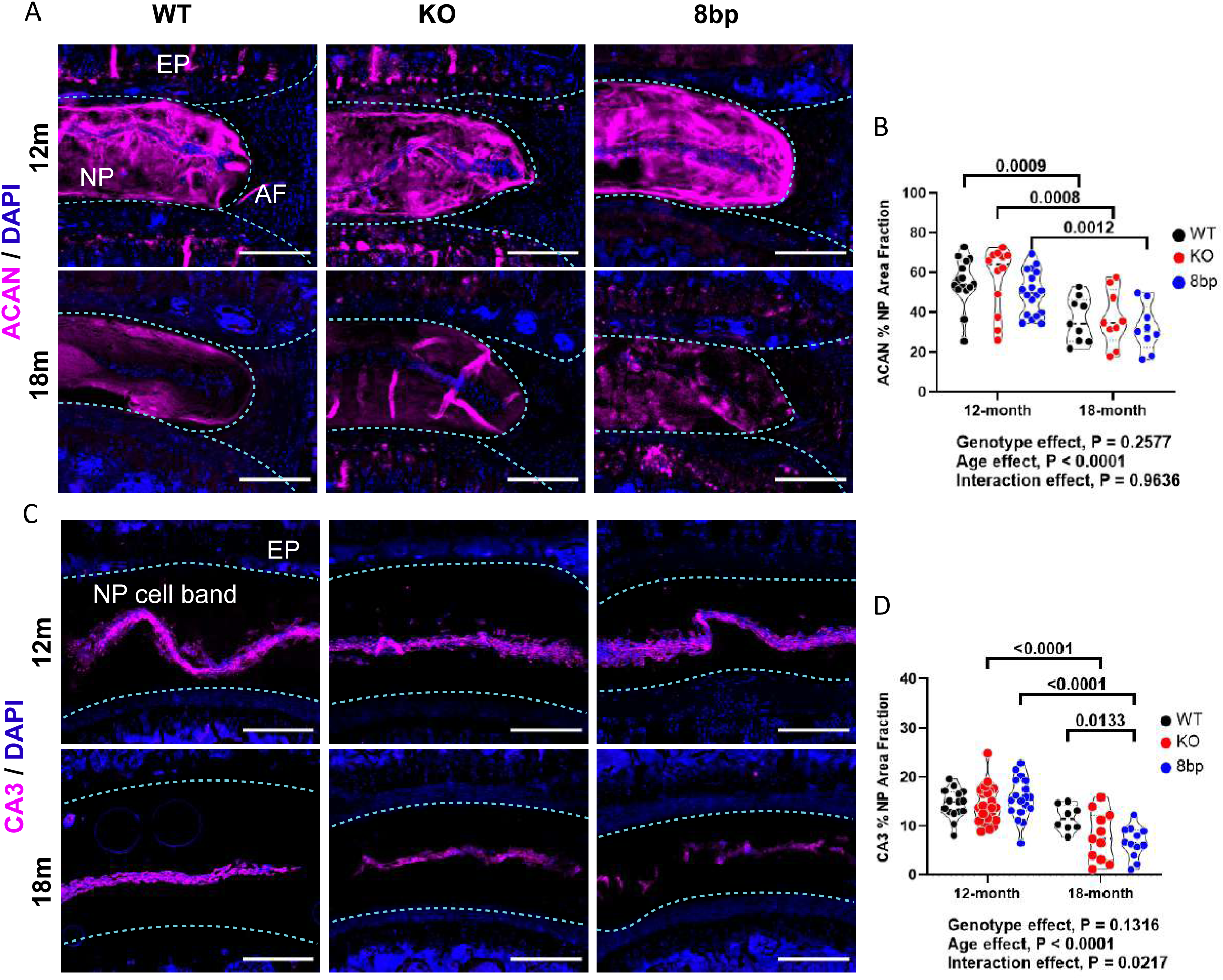
*Chadl* loss results in changes in NP cell phenotype. (A) Representative images showing Aggrecan staining of 12M and 18M disc sections from WT, KO, and 8bp mice. (B) Percent area quantification of Aggrecan in NP compartment. (C) CA3 staining of 12M and 18M disc sections of WT, KO, and 8bp. (D) Percent area quantification of CA3 in NP compartment. Violin plots show score distribution with median and quartile range, *P* < 0.05. For 12M: 5-8 WT, KO, 8bp mice, n = 2-4 discs/mouse, 14-24 discs total per genotype. For 18M: 3-6 WT, KO, 8bp mice, n = 2-4 discs/mouse, 9-14 discs total per genotype. Significance was determined using 2-way ANOVA with Tukey’s Post Hoc Test, comparing Age and Gene interaction. Scale bar 250µm.

### CHADL loss causes changes in vertebral bone

Previous work has implicated CHADL as a regulator of chondrocyte differentiation (15). Accordingly, to assess the impact of CHADL loss on vertebral bone during aging, we performed µCT of the lumbar vertebrae (L1-6) in mice at 12 and 18 months. Three-dimensional (3D) reconstruction of vertebra trabecular tissue highlighted bone loss during the natural aging in wildtype (WT) mice, but CHADL loss affected trabecular bone architecture in middle-aged and old mice (Fig. 7A, B). µCT analysis revealed that in 18-month-old *Chadl* mutant mice compared to WT mice, there was a small increase in percent bone volume (BV/TV), trabecular number (Tb.N.), with a concomitant decrease in trabecular thickness (Tb.Th.) and separation (Tb.Sp.) (Fig. 7C-F). There were no significant differences in trabecular bone mineral density between the genotypes at either time point (Fig. 7G). As in trabecular tissue, the cortical lumbar vertebrae were also affected in *Chadl* mutant mice (Fig. 7H-M). It was notable that only *Chadl* mutants showed a decrease in cortical bone volume (BV) as they aged (Fig. 7H). The 8bp mice showed a reduction in cross-sectional thickness (Cs. Th.) at both ages and a slight increase in Bone Perimeter (B.Pm) at 18 months without notable differences in mean total cross-sectional bone area (B.Ar.) and tissue area (T.Ar.) (Fig. 7I-L). Moreover, a significant reduction in tissue mineral density (TMD) was noted in both *Chadl* mutant mice at 18 months (Fig. 7M). These changes in vertebral bone health parameters suggest that CHADL plays an important role in regulating trabecular and cortical vertebral bone geometry.

**Figure 7.**
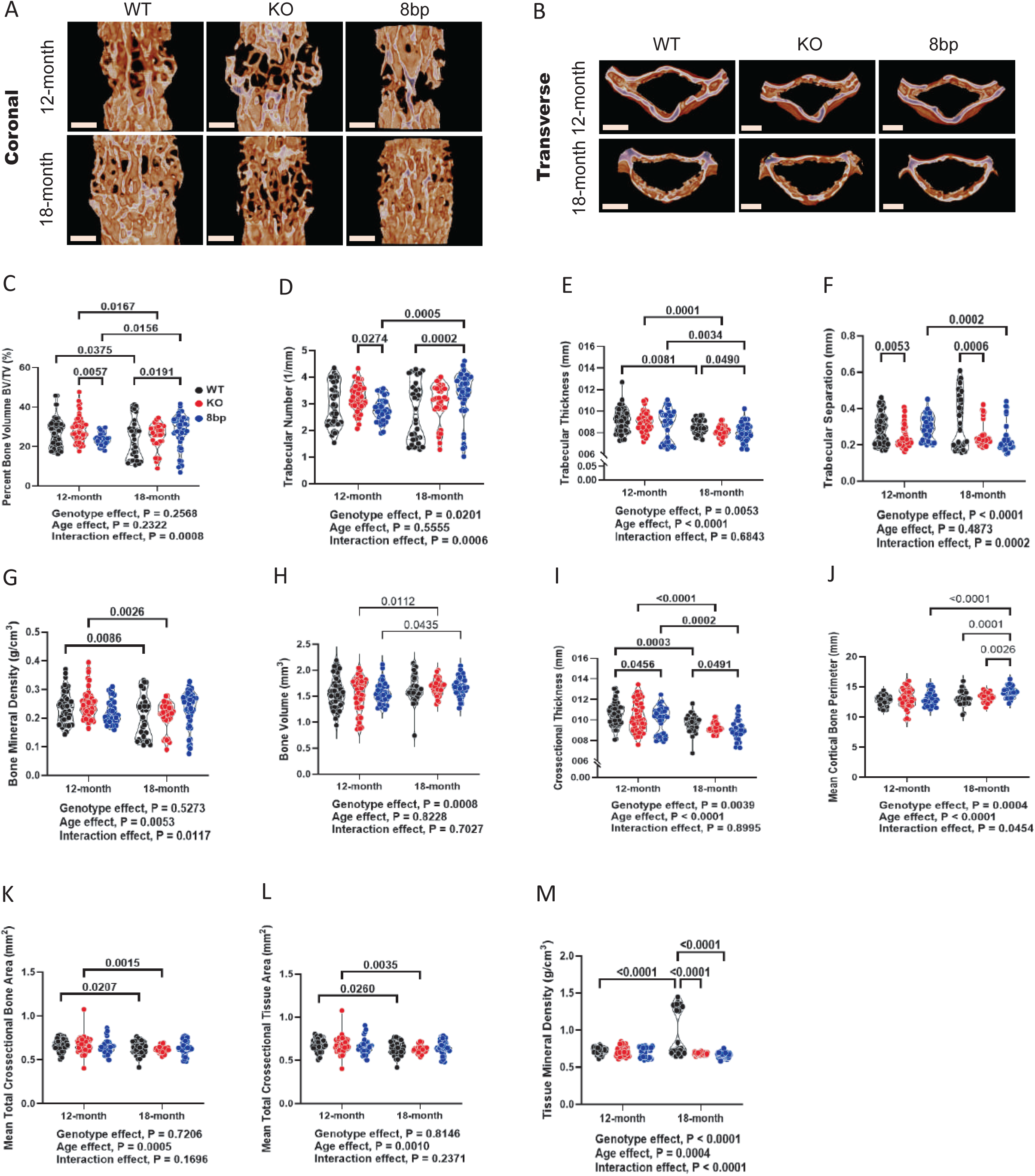
Chadl loss contributes to altered vertebral trabecular and cortical bone parameters. (A) Representative coronal images of vertebral trabecular bones of WT, KO, and 8bp mice. (B) Transverse image of the cortical shell of vertebrae from WT, KO, and 8bp mice. Vertebral trabecular structural parameters (C) Percent bone volume BV/TV (%), (D) Trabecular Number (1/mm), (E) Trabecular Thickness (mm), (F) Trabecular Separation (mm) and (G) Bone mineral density (g/cm^3^). Cortical bone structural parameters (H) Bone Volume (mm^3^), (I) Cross-sectional Thickness (mm), (J) Mean cortical bone perimeter (mm), (K) Mean total cross-sectional bone area (mm^2^), (L) Mean total cross-sectional tissue area (mm^2^), (M) Tissue mineral density (g/cm^3^). Violin plots show score distribution with median and quartile range, *P* < 0.05. For 12M: WT (8F, 3M), n = 66 discs and vertebral bones total; KO (6F, 4M), n = 48; 8bp (5F, 2M), n = 42. For 18M: WT (3F, 3M), n = 36 discs and vertebral bones total; KO (1F, 5M), n = 36; 8bp (1F, 5M), n = 42). Significance was determined using 2-way ANOVA with Tukey’s Post Hoc Test comparing Age and Gene interaction. Scale bar 500µm.

## Discussion

In this study, we employed two independent mouse models of CHADL loss-of-function, a global null allele and a human-informed frameshift mutation analogous to the osteoarthritis (OA)-associated variant (13), to systematically examine the consequences of deficiency across skeletal tissues during aging. Our findings demonstrate that loss of CHADL results in age-dependent degeneration of the intervertebral disc, with a smaller impact on knee articular cartilage OA incidence, and minimal impact on hip articular cartilage health, despite *CHADL* being one of the strongest known genetic risk factors for human hip OA (13). These results uncover a previously unrecognized tissue-selective role for CHADL in maintaining spinal disc extracellular matrix (ECM) homeostasis, while providing important mechanistic insight into the joint-specific nature of OA genetic risk.

A central and unexpected finding of our work is that CHADL deficiency preferentially affects the lumbar intervertebral disc (IVD) rather than other synovial joints. Both *Chadl* mutant lines developed progressive disc degeneration with aging, characterized by altered disc height and disc height index, and disruption of annulus fibrosus (AF) collagen architecture, evidenced from abnormal proportions of thin and intermediate thickness collagen fibrils observed under polarized light, indicating dysregulated collagen turnover rather than a simple reduction in matrix content. These findings suggested that the degenerative cascade was characterized by changes in the ECM’s water-binding properties and by overall tissue swelling (26). These conclusions were supported by reduced abundance of key fibrous ECM components (including COL1, COL2, COMP, and FMOD), and loss of nucleus pulposus (NP) phenotypic identity as evidenced by lower CA3 expression. These changes are highly consistent with CHADL’s established role as a non-glycanated small leucine-rich proteoglycan (SLRP) that binds fibrillar collagens and modulates ECM organization (15). The intervertebral disc, particularly the AF, is a tissue uniquely dependent on precise collagen fibril assembly, alignment, and turnover to withstand complex tensile and compressive forces (27). Unlike articular cartilage, which relies more heavily on aggrecan-mediated osmotic swelling pressure (28), the AF is predominantly a collagen-rich, load-bearing structure, making it especially vulnerable to perturbations in collagen-binding regulatory proteins such as CHADL. Thus, our findings suggest that CHADL functions as a molecule that fine-tunes disc ECM architecture, with its loss rendering the disc particularly susceptible to age-related degeneration.

In contrast to the disc phenotype, CHADL loss exerted a small effect on knee OA in middle-aged mice but did not exacerbate OA in the hip and knee joints even in aged mice. Histopathological evaluation and OARSI scoring confirmed comparable age-related OA progression across genotypes in aged mice. These findings mirror the joint-specific effects observed in large-scale human genetic studies, where *CHADL* variants show very strong associations with hip OA, modest or inconsistent effects at the knee, and minimal impact at other joints (13).

The lack of overt hip OA in *Chadl* mutant mice underscores several important concepts. First, genetic risk does not translate uniformly across joint sites, even for variants with exceptionally large effects in humans. Second, CHADL’s role in ECM regulation appears to be highly context-dependent, influenced by differences in tissue architecture, mechanical loading, and compensatory matrix networks. Articular cartilage expresses multiple glycanated and non-glycanated SLRPs (such as chondroadherin, decorin, biglycan, fibromodulin, and lumican) that may functionally compensate for CHADL loss in joints but not in the disc. Third, the strong association of *CHADL* with human hip OA may reflect long-term, biomechanical, or developmental processes that are not fully recapitulated in murine hip joints, which differ substantially from humans in size, loading patterns, lifespan, and gait.

Human genetic studies identify a frameshift variant in *CHADL* (rs532464664) that confers one of the largest known effect sizes for hip OA, operates through a recessive mechanism, and leads to nonsense-mediated decay of the full-length transcript (13). A subsequent analysis confirmed the OA risk associated with the frameshift but in this case, it was indicated by the proxy SNP rs117018441 (14). The risk T allele at rs117018441 is highly correlated to the frameshift variant (r^2^ = 0.8) and therefore despite the association of the T allele with higher *CHADL* transcript in various tissues of the Genotype-Tissue Expression (GTEx) Portal, it is unlikely that these transcripts produce functional protein. Of the OA risk genes analyzed, Tuerlings et al. identified *CHADL* as one where expression was increased in lesional vs. preserved cartilage and subchondral bone from end-stage OA (29). Functional data indicate that CHADL acts as a negative regulator of chondrocyte differentiation and a collagen-binding ECM protein (15). Together, the human genetic and transcriptomic evidence and our mouse studies indicate that CHADL likely plays a role in matrix production of multiple musculoskeletal tissues and could be a pathway worth investigating for optimizing tissue homeostasis.

Our *in vivo* data suggest that one major consequence of CHADL loss is destabilization of collagen-rich tissues subject to complex mechanical demands, particularly the intervertebral disc. Notably, independent transcriptomic studies of OA have identified *CHADL* as one of the most consistently differentially expressed genes in both cartilage and subchondral bone, alongside *IL11* (29). Moreover, recent mechanobiology studies demonstrate that *CHADL* expression is strongly mechanosensitive, showing marked down-regulation under high hydrostatic pressure (30). Given that the disc experiences sustained compressive and hydrostatic loads throughout life, dysregulation of CHADL-mediated mechanoadaptation may represent a critical vulnerability in this tissue.

These observations raise the possibility that human *CHADL*-associated hip OA may involve coupled degeneration of the hip joint and adjacent spinal structures, or that disc pathology may contribute indirectly to altered biomechanics and hip joint loading over time. Importantly, disc degeneration was not evaluated in most human OA genetic cohorts, suggesting that spinal pathology may be an underappreciated component of CHADL-mediated disease risk.

In addition to disc degeneration, CHADL loss altered vertebral trabecular and cortical bone architecture, particularly in aged mice. Changes in trabecular organization, cortical thickness, perimeter and tissue mineral density indicate that CHADL influences not only cartilaginous tissues but also vertebral bone remodeling, consistent with its role in regulating chondrocyte differentiation and matrix assembly during endochondral processes. These skeletal effects further support the concept that CHADL is a spine-relevant ECM regulator, acting at the interface of disc, cartilage, and bone.

Collectively, our findings redefine the functional landscape of CHADL in the adult skeleton, identifying the intervertebral disc as a primary tissue susceptible to CHADL deficiency. This work emphasizes the importance of tissue- and joint-specific interpretation of OA genetic risk and cautions against assuming uniform effects across synovial joints. From a translational perspective, CHADL emerges as a potential therapeutic target for spinal degeneration, while its role in joint OA appears to be more nuanced, context-dependent, and perhaps dependent on co-variants in other loci.

Future studies should address whether disc pathology is present in human carriers of CHADL risk variants, explore interactions between CHADL and other SLRPs in cartilage and disc, and determine how CHADL-mediated mechanosensing influences long-term tissue adaptation. Such efforts will be essential for leveraging genetic discoveries into targeted interventions for both OA and degenerative spine disease.

## Materials and Methods

### Sex as a biological variable

Our investigation examined male and female mice, and similar findings are reported for both sexes.

### Generation and validation of *Chadl* mouse lines

All animal care procedures, housing, breeding, and the collection of animal tissues were performed in accordance with a protocol approved by the Institutional Animal Care and Use Committee (IACUC) of the University of North Carolina at Chapel Hill. The *Chadl* knockout allele was provided by the International Mouse Phenotyping Consortium (IMPC) and was generated at the Wellcome Trust Sanger Institute. The allele name is *Chadl^em1(IMPC)Wtsi^* with Mouse Genome Informatics (MGI) identification number 6153306. The knockout was generated via 897 bp deletion (37 base pairs upstream of exon 3 plus the first 860 base pairs of exon 3) that was mediated by guide RNAs 5’-CCACTCGACCAGGGGCCGTGCGT-3’ and 5’-CCATTAGTTCATAACACCACCGG-3’. The genotyping primers of common forward (5’-CCTTTTCGTGTGGCTGAGAT-3’), wild type reverse (5’-GTTGTGGGCCAAGTTGAGAG-3’), and knockout reverse (5’-GGGCAGCGCAGGTCAGAG-3’) yield products of 582 base pairs for the wild type allele and 255 base pairs for the KO allele. The *Chadl* 8 bp allele was generated by the Animal Models Core at the University of North Carolina at Chapel Hill. Pronuclear microinjection of zygotes was used for genome modification, with the injection mix containing two ribonucleoprotein (RNP) complexes of guide RNAs and Cas9 (guide 1: GGTGCGCGCTCGGGTGCGCT ; guide 2: CGCGCTCGGGTGCGCTCGGA) and a 113 bp single-stranded oligonucleotide donor that contains the 8 base pair insertion (bolded) and is antisense to the Chadl coding region: CGCAGGCGCCGCGGGCCCCGGCACGCGCCATCCGAGCGCA**CGCGCGCC**CCCGA GCGCGCACCAGCCACTCGAGCAGGGGCCGTGCGTGGCAGGCGCACCACAGCGGGTTGCCC. After injection, embryos were transferred to pseudopregnant females for development. Founder pups were screened by genotyping and Sanger sequencing to detect successful insertion. Progeny were genotyped by a common primer set of Forward (5’-CTGCTCGAGTGGCTGGTG-3’) and Reverse (5’-GGGCAGCGCAGGTCAGAG-3’, same as knockout reverse). Products run on 4% agarose for extended period of time can be distinguished for the 113 bp (wild type) and 121 bp (8 bp insertion) amplicons.

### Western blot for confirmation of CHADL knockout

Hip cartilage explants from 2-week old mice were obtained from wild type, KO, and 8 bp mice. Protein was isolated using 500 µl of TRIzol in bead beater tubes (VWR 10158-610) with two 45-second runs of 6500 on a Precellys 24^®^ (Bertin Corp). Phase separation by 12,000g for 10 minutes at 4 °C generated an insoluble ECM fraction that was retained and combined with the phenol ethanol layer for protein isolation.

Isopropranol precipitation was followed by washes with 95% and then 100% ethanol. The protein was then solubilized with a buffer containing 20 mM EDTA, 140 mM NaCl, 5 % SDS, and 100 mM Tris, with supplementation by PMSF and Halt protease inhibitor cocktail (31). Solubilization was performed at 50 °C for 24 hours under mixing by ThermoMixer C at 400 rpm. Protein quantification was performed by Bradford assay and then 15 µg was separated by SDS-PAGE and transferred to nitrocellulose membrane. The membrane was incubated with CHADL antibody (HPA024654, Sigma-Aldrich, 1:500) overnight, followed by a secondary antibody, and chemiluminescent images were obtained using the Azure c600 imaging system. The membrane was stripped, cut, and incubated with antibodies against two control proteins: Collagen VI (sc-377143, Santa Cruz, 1:50), and GAPDH (2275-PC, Trevigen, 1:2000).

### Histological Analysis

Mice were sacrificed by CO_2_ euthanasia with secondary physical method. Hindlimbs were dissected and the femur was bisected to establish samples for the knee (stifle) and hip (with pelvis included). As with prior studies, joints were fixed for 4 days in 4% paraformaldehyde (PFA) at 4 °C, decalcified in Immunocal for 4 days at room temperature, and then processed for paraffin embedding by the UNC Pathology Services Core Facility (32). Both knee (stifle) and hip joints were embedded for coronal sectioning. Mid-coronal sections of the knees were stained with Safranin O / Fast Green / Hematoxylin and all four quadrants (lateral femur, lateral tibia, medial femur, medial tibia) were evaluated on the 0-6 OARSI scale (max total joint score = 24) (25). Coronal sections of hip joints (n=10 wild type, n=6 KO, n=6 8bp) were stained with Safranin O / Fast Green / Hematoxylin and evaluated for cartilage degradation.

Lumbar spines were also dissected and immediately fixed in 4% PFA in PBS at 4°C for 48 hours, decalcified in 20% EDTA at 4°C for 2-3 weeks, and then embedded in paraffin. 7 µm mid-coronal sections from L1-S1 levels were deparaffinized, stained with Safranin-O/Fast Green/Hematoxylin, and images acquired on a light microscope (Axio Imager 2; Carl Zeiss Microscopy) using 5x/0.15 N-Achroplan (Carl Zeiss) objective and Zen2™ software (Carl Zeiss). The health of disc compartments was assessed by at least four blinded graders using Modified Thompson Grading (33-35). Picrosirius red staining (Polysciences, 24901) was performed to assess collagen fibril thickness, and images were acquired using 4x /0.25 Pol /WD 7.0 (Nikon) objective on a polarizing light microscope (Eclipse LV100 POL; Nikon) (36, 37). NIS Elements Viewer software (Nikon) was used to set the color threshold for the level of green (thin), yellow (intermediate), and red (thick) fiber. Color threshold levels remained constant for all samples.

### Micro-Computed Tomography (µCT) Analysis

*Chadl* 8-bp, KO, and WT lumbar spines were dissected and fixed in 4% PFA in PBS at 4°C for 48 hours prior to µCT scans (Bruker, Skyscan 1275). For assessment of bone morphometry, the lumbar spines of *Chadl* KO, 8bp and WT mice were rinsed and hydrated in 1x PBS, scanned at a resolution of 15 µm^3^ voxel (50 kV, 200 µA, 85 ms exposure time, rotation step of 0.2°, using 1 mm aluminum filter). The reconstruction was performed using Skyscan NRecon package. Distance measurements were made along the dorsal, midline, and ventral regions in the sagittal plane of disc height and vertebral length and then averaged, which was then used to calculate the disc height index (DHI), as previously described (34, 35). Trabecular parameters were measured using Skyscan CT analysis (CTAn) software by contouring the region of interest (ROI) in the 3D reconstructed trabecular tissue. Resulting datasets were assessed for bone volume fraction (BV/TV), trabecular number (Tb. N.), trabecular thickness (Tb. Th.), and trabecular separation (Tb. Sp.). The cortical bone was analyzed in two dimensions and assessed for bone volume (BV), cross-sectional thickness (Cs. Th.), mean cross-sectional bone area (B. Ar), mean cross-sectional tissue area (T. Ar) and mean cortical bone perimeter (B. Pm.). Mineral density was calculated using a standard curve created with a mineral density calibration phantom pair (0.25 g/cm^3^ calcium hydroxyapatite (CaHA) and 0.75 g/cm^3^ CaHA).

### Immunohistochemistry and digital image analysis

Mid-coronal disc sections (7 μm) were deparaffinized in histoclear and rehydrated in ethanol solutions (100–95%), water, and PBS. Citrate-buffer at pH 6 (Vector Laboratories) antigen retrieval method was performed on samples required for COLI (1:100; Abcam, ab34710), COLII (1:400; Fitzgerald, 70R-CR008), CA3 (1:150; Santa Cruz Biotechnology, sc-50715); FMOD (1:250; Abcam, ab81443); 1:1000 Proteinase K antigen retrieval for COMP (cartilage oligomeric matrix protein) (1:200; Abcam, ab231977); 1:200 chondroitinase ABC (20 U/mL) enzymatic retrieval for ACAN (1:50; Millipore Sigma, AB1031). MOM kit (Vector Laboratories; BMK-2202) was used per manufacturer’s instructions. After antigen retrieval, samples were blocked with 5-10% normal goat or donkey serum (Jackson ImmunoResearch) in PBS-Triton (0.4% Triton X-100 in 1x PBS) for 1 h at room temperature. Primary antibodies were applied and incubated overnight at 4°C. Samples were then generously washed three times with PBS and incubated with Alexa Fluor-594-conjugated secondary antibody (1:700, Jackson ImmunoResearch) for 1 h at room temperature, shielded from light. Samples were washed three times with PBS and mounted with ProLong Gold Antifade Mountant with DAPI (Thermo Fisher Scientific, P36934). Images were acquired on Axio Imager 2 (Carl Zeiss Microscopy) using 5×/0.15 N-Achroplan (Carl Zeiss Microscopy) or 10×/0.3 EC Plan-Neofluar (Carl Zeiss Microscopy) objective, X-Cite 120Q Excitation Light Source (Excelitas Technologies), AxioCam MRm R3 camera (Carl Zeiss Microscopy), and Zen 2^TM^ software (Carl Zeiss Microscopy). Per staining experiment, images were taken at a set exposure time for all samples. All quantifications were done on ImageJ 1.52i (NIH). Images were thresholded to create binary images, and NP and AF compartments were manually contoured using the Freehand Tool.

### Statistical analysis

Statistical analysis was performed using Prism 10 (GraphPad, La Jolla, CA, USA), and data were presented as violin plots showing all data points, with median, interquartile range, and maximum and minimum values. Differences between distributions of two samples were checked for normality using the Shapiro-Wilk test. One-way ANOVA with Sidak’s post hoc test or a Two-Way ANOVA with Tukey’s post hoc test was used to determine significance between groups and the interaction between genotype and age. Since interactions between genetic, biological, and biomechanical factors at individual spinal levels have been shown to produce different phenotypic outcomes, each intervertebral disc or vertebra is considered an independent sample, a methodology followed by many groups (33-35). Analyses of Modified Thompson Grading data distributions and fiber thickness distributions were performed using a chi-square test at a 0.05 level of significance.

## Data availability

All the underlying data and supporting information for the article is included in the manuscript.

## Author Contributions

BOD, MVR designed research studies. KS, BM, JS, AM, EVF, JRF, CDS, TMO, SD, WAC conducted experiments, acquired and analyzed data. KS, BOD and MVR wrote the manuscript.

## Acknowledgements

We thank the University of North Carolina Animal Models Core for technical assistance in generating *Chadl* 8bp mice. We thank Sarah E. Bell for helping with the manuscript preparation and schematics.

