## Supplementary Figure for "Human hip osteoarthritis–associated *Chadl* variant induces intervertebral disc degeneration in mice"

SFig1.

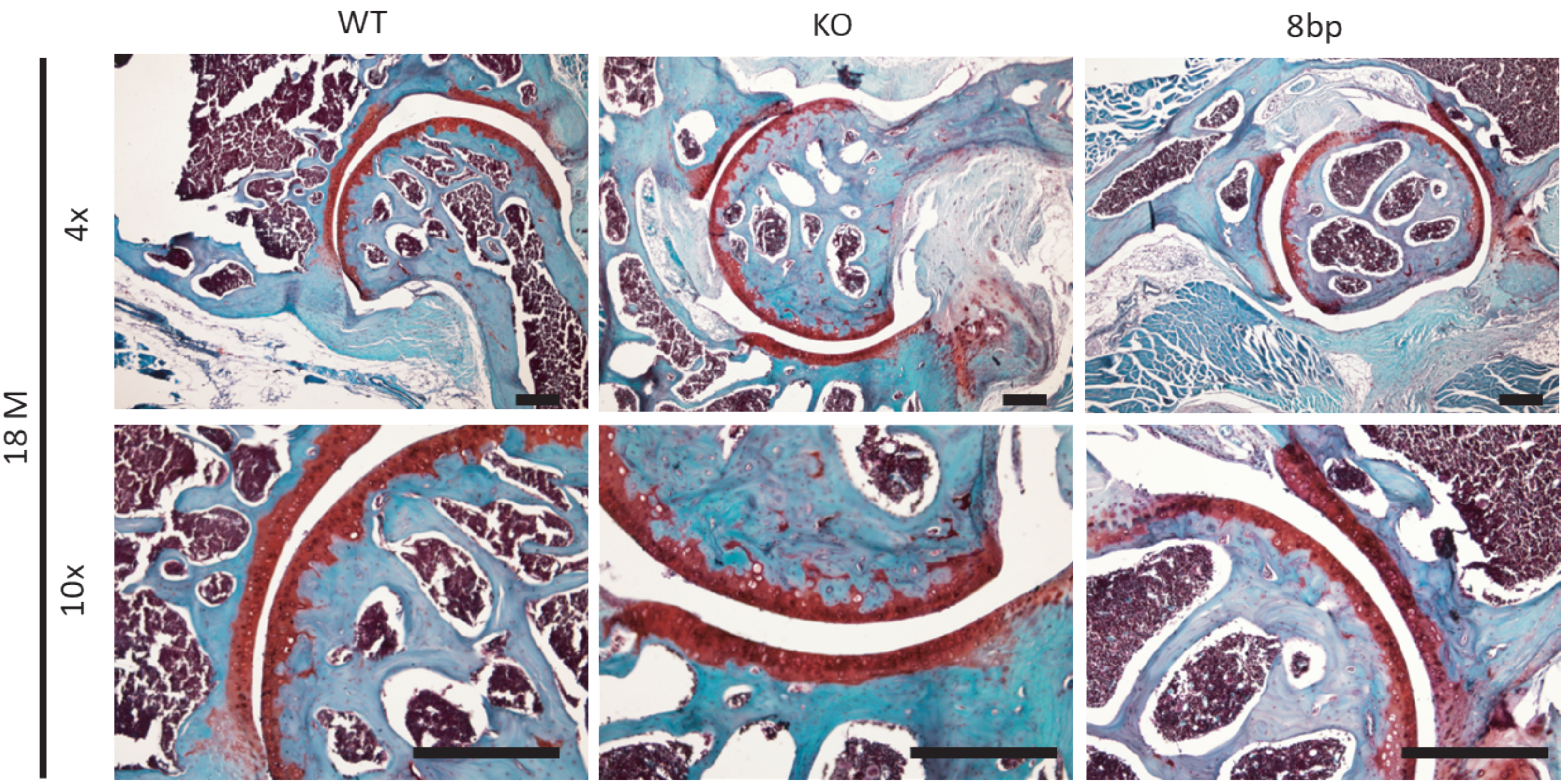


Supplementary Figure 1. (A) Representative pictures of synovial hip joints of aged mice (≥18 months old) at 4X and 10X magnification from wildtype (n=10), knockout (n=6), and 8bp (n=6) mice.
